# MetaGXplore: Integrating Multi-Omics Data with Graph Convolutional Networks for Pan-cancer Patient Metastasis Identification

**DOI:** 10.1101/2024.06.30.601445

**Authors:** Tao Jiang, Haiyang Jiang, Xinyi Ma, Minghao Xu, Yan Liang, Wentao Zhang

## Abstract

The spread of cancer cells from the primary tumor to distant anatomical sites, known as tumor metastasis, poses a significant challenge in clinical prognosis, impairing treatment efficacy and reducing patient survival time. Current methods for predicting and diagnosing tumor metastasis rely heavily on extensive examinations and subjective clinical judgments. Accurate and rapid prediction of tumor metastasis likelihood remains an unresolved challenge, crucial for guiding effective clinical interventions and extending patient survival. Additionally, identifying key genes highly associated with metastasis probability is a pressing issue, essential for providing valuable insights into the potential identification of tumor metastasis-specific biomarkers. We developed MetaGXplore, a pioneering Graph Convolutional Neural Network (GCN)-based framework designed to predict metastasis probability by integrating pan-cancer multi-omic datasets with a protein-protein interaction network, while also identifying key genes involved in the metastatic process. Multi-omics datasets offer a comprehensive view of cancer biology, enhancing accuracy in metastasis forecasting through superior deep learning algorithms. Our classification model results were interpreted with GNNExplainer. The efficacy of MetaGXplore was validated via model evaluations, graph structure analysis, and multi-omics data assessment. Enrichment analysis of key genes further explored their biological roles.

## I. Introduction

Cancer metastasis, a critical hallmark of malignancy associated with tumor progression and poor patient prognosis, involves the dissemination of cancer cells from the primary tumor to distant organs. Metastasis presents a formidable challenge in cancer management, often signifying advanced tumor progression and adversely affecting patient survival outcomes. Conventional methods for identifying tumor metastasis primarily rely on histopathological examinations, imaging techniques, and clinical discoveries. While these methods provide relatively accurate assessments of overt metastasis, they depend heavily on the expertise of healthcare professionals and may lack sensitivity in detecting micrometastasis or accurately predicting metastatic risk [1, 2, 3, 4, 5, 6, 7]. Consequently, there is an urgent need for innovative tools that offer convenient and rapid assessment of tumor metastatic risk to aid clinical management [8, 9, 10]. In recent years, leveraging artificial intelligence (AI), particularly deep learning methods, has emerged as a transformative approach in disease research and clinical practice. AI-based predictive models utilizing multi-omics data have shown significant potential across various applications. The advancement of high-throughput sequencing technology and the advent of multi-omics sequencing have exponentially increased the molecular characteristics that can be used as features and insights into pathophysiology [11, 12, 13, 14]. However, the inherent heterogeneity of cancer results in diverse molecular characteristics among patients. This patient-specific molecular signature can be conceptualized as a unique subnetwork within the broader molecular network, representing individual genetic features [3]. Integrating and transforming these patient genetic features into patient-specific networks allows for deeper insights, improving the detection of tumor metastasis risk through AI-driven analysis and enhancing prognostic treatment strategies. Current approaches for predicting cancer metastasis using machine learning face significant challenges, such as the high dimensionality of data points representing individual patient features from multi-omics datasets [15]. This “curse of dimensionality” leads to instability in the feature selection process across different datasets. Recently, graph convolutional neural network (GCN) has been widely adopted for processing graph signal classification tasks, effectively addressing the “curse of dimensionality” [16, 17]. Furthermore, interpreting deep learning model predictions within a genetic context is essential, especially given regulatory requirements like the European Union’s General Data Protection Regulation (GDPR), which imposes restrictions on automated decisions generated by algorithms [18]. The interpretability of deep neural networks is thus a critical factor in enhancing confidence in AI model predictions and understanding the mechanisms underlying tumor metastasis. In this study, we introduce MetaGXplore, a novel deep learning model that integrates Graph Convolutional Networks (GCNs) with multi-omics pan-cancer data to predict metastasis. Our model was trained and tested on a dataset comprising 754 samples from 11 cancer types, each with balanced evidence of metastasis and non-metastasis. The results demonstrate MetaGXplore’s superior performance compared to other deep learning methods such as Multi-Layer Perceptron (MLP) and transformers. Additionally, to enhance MetaGXplore’s interpretability, we employed Graph Mask and Feature Mask methods from GNNExplainer. These techniques elucidated the model’s inner workings and identified key genes associated with metastasis. Subsequently, pathway enrichment analysis provided insights into the functions of these genes and explored their interactions within Protein-Protein Interaction (PPI) networks, supplemented by literature reviews to comprehensively interpret their roles. Overall, our study advances the application of AI and multi-omics data in clinical treatment strategy selection and provides a valuable gene set for future research on tumor metastasis.

The main contributions of the study are summarized as follows:

- We present the first application of Graph Convolutional Networks (GCNs) to integrate pan-cancer multi-omics data for accurate prediction of tumor metastasis.
- A novel interpretability approach was introduced using Graph Mask and Feature Mask methods from GNNExplainer, allowing key metastasis-associated genes to be effectively identified.
- Extensive experiments conducted on a dataset of 754 samples from 11 cancer types demonstrate that MetaGXplore significantly outperforms other deep learning models, including Multi-Layer Perceptron (MLP) and transformers.

## II. MATERIALS AND METHODS

In this part, we describe the data preparation process, the construction of the patient multi-omics network, the framework of MetaGXplore model, and the implementation of the explainable model, as illustrated in Fig.1.

**Fig. 1:**
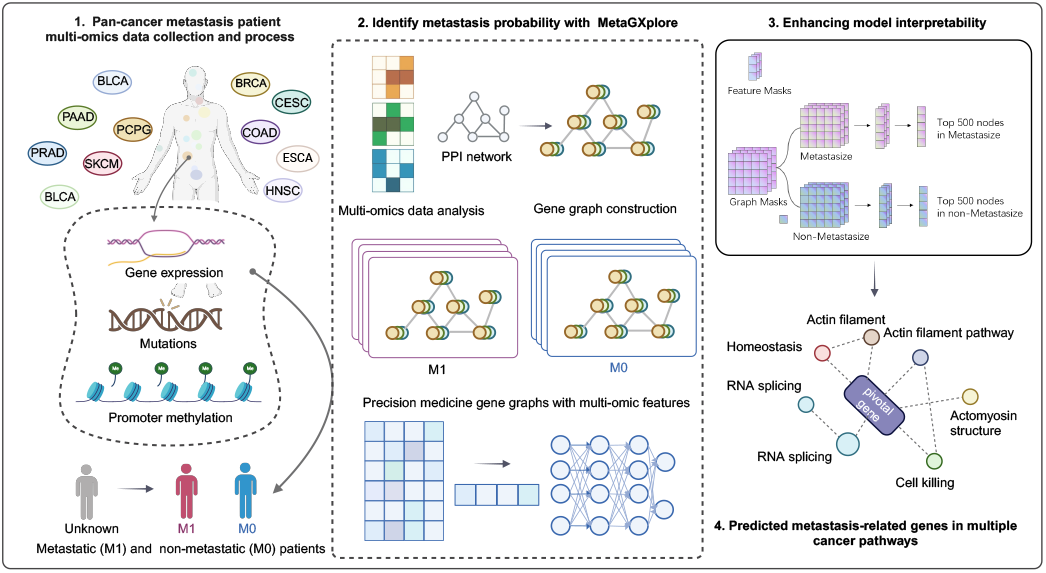
Schematic overview of MetaGXplore and explainable approach for identifying metastasis-related genes.

### A. Data Collection and Preprocessing

We collected primary samples from 33 different cancers from The Cancer Genome Atlas (TCGA) projects, each with and without distant metastasis. The dataset includes mRNA, methylation, and mutation data. Specifically, we focused on 11 cancer types where all three types of omics data are available simultaneously. Among the datasets, 754 samples were used for modeling, comprising 377 samples diagnosed with metastatic cancer and an equal number of 377 samples diagnosed with primary tumors without evidence of metastasis. The categorization of distant metastasis status follows the American Joint Committee on Cancer (AJCC) guidelines, with M0 assigned to samples without evidence of distant metastasis and M1 to those that show such evidence (4). Equal counts are evident for both metastatic and in situ cancers, as depicted in Table I.

**TABLE I:**
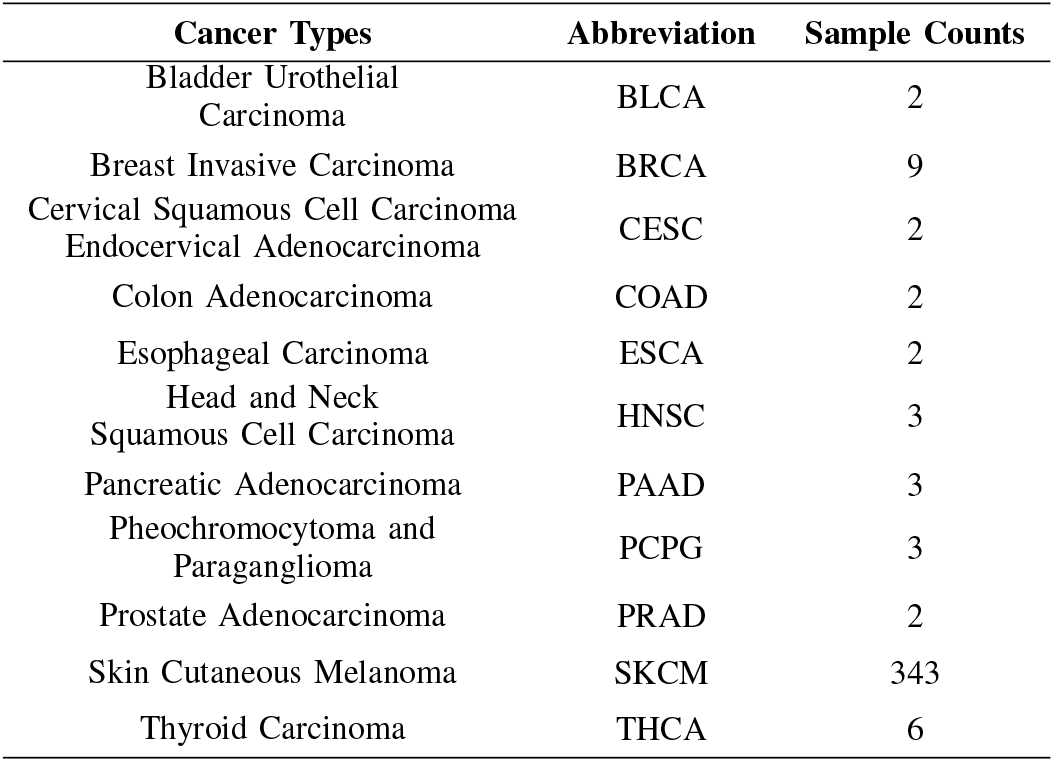
DISTRIBUTION OF CANCER TUMOR TYPES AND SAMPLE COUNTS IN METASTATIC.

The multi-omic datasets were acquired using the TCGA-biolinks R package, enabling access to the Genomic Data Commons (GDC) Data Portal (2; 8). Utilizing both the TCGA-biolinks and TCGA-assembler packages, we conducted preprocessing on the harmonized TCGA dataset. Initially, we mapped CpG islands within 1500 bp upstream of gene transcription start sites (TSS) and computed average methylation values. To address missing values, we implemented three preprocessing steps based on a reported approach (1). First, we excluded genes (e.g., mRNA/DNA methylation) with zero values in more than 25% of patients. Next, we filtered out samples with less than 75% of the remaining features. For mutation data, we excluded features with zero occurrences across all samples and filtered out features with zeros in more than 40% of patients. Finally, we employed the R impute function (7) to impute missing expression data using nearest neighbor averaging to fill the remaining gaps. Each test set underwent independent preprocessing. After preprocessing, there were 377 metastatic samples encompassing all three data types. For interpretability purposes, we retained genes featuring all three omics data types and randomly selected an equivalent number of non-metastatic samples for statistical analysis.

For the network-based approach, we downloaded the PPI information from Database STRING v12.0 (https://string-db.org/). We merged this information to construct a huge network and then mapped the genes with multi-omics features onto the network to obtain a comprehensive interacted multi-omic gene network. To transform omics data into a graph format, each patient can be represented as a graph: 𝒢= 𝒢 _1_, *𝒢* _2_, …, *𝒢* _*N*_, where N is the number of patients. For an arbitrary graph 𝒢_*i*_ = (𝒱 _*i*_, ℰ_*i*_, **X**_*i*_,**Y**_*i*_), 𝒱_*i*_ and ℰ_*i*_ denote the nodes and edges, which represent genes and protein-protein interactions, respectively. 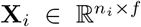 is the node feature matrix, where |𝒱| _*i*_ = *n*_*i*_ is the number of genes, and *f* is the number of omics features. **Y**_*i*_ is the label of the ith graph, with **Y**_*i*_ = 1 if the ith patient has metastasized and **Y**_*i*_ = 0 otherwise. Constructed by 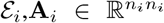 is the adjacency matrix describing the graph’s edge connection information.

### B. Node Embedding Extractor

#### a) 1) Multi-head attention mechanism in Transformer

Owing to its outstanding expressive ability, the multi-head attention mechanism is the most crucial component in transformers for extracting embeddings from text or image data. Recently, some research has applied this model to graph data problems. The vanilla transformer [19] calculates pairs of attention weights among all data entities and updates the embeddings according to the weights from all other data entities. However, this type of transformer model does not account for graph structure information. To capture topological information, the graph transformer [20] calculates attention only between nodes and their neighbors:

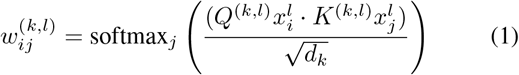

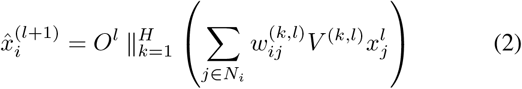

In Eq.1, in lth layer, 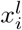 and 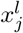 represent features of current node *i* and neighbor nodes of *j*. The *k*th weight 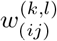 determines the importance of node *i* and neighbor *j* and is obtained by node vector dot product after the linear transformed of learnable parameters (Q,K), and this weight is normalized by dimension of node features. In Eq. 2, H means parallel attention mechanisms with different parameter initialization to improve the stability and generalization ability when train the model. Neighbor features that are linear transformed by trainable weight *V* ^(*k,l*)^ are weighted summed and H heads are concatenated. Dimension of this result is H times than the input feature and the learnable matrix *O*^*l*^ are used to map this high dimension feature matrix to the same dimension as the input feature. As for vanilla transformer, node *j* means nodes other than node *i*.

#### b) Graph convolution network

GCN [21] is a graph model to extract embedding from graph data based on spectral clustering. This model needs two inputs: adjacency matrix **A** ∈∈ ℝ^*n×n*^ and multi-omics feature matrix **X** ∈∈ ℝ^*n×d*^. GCN can be built by stacking many layers, lth layer is defined as:

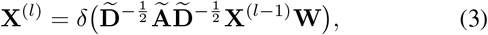

Here, 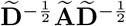 denotes the normalized graph Laplacian, **Ã** = **A** + **I**_*n*_ denotes adjacency matrix with added self-connections, 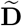 is degree matrix of **Ã**, **W** is trainable weight matrix. *δ* denotes the nonlinear activation function, generally the ReLU activation function, and **H**^(*l*)^ is the input of each layer, and notably, **H**^(0)^ = **X**.

### C. Figures and Tables Prediction of Patient Metastasis Classification

In the classification model, we stack two node embedding extraction layers, such as GCN, fully connected layers, transformer, and graph transformer layers, each followed by a ReLU activation function. After the embedding layers, to obtain a vector containing information about the entire graph, we use a readout layer [22, 23] to aggregate node embeddings:

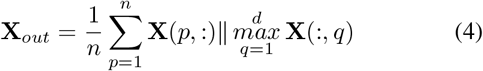

Finally, we use a Multilayer Perceptron (MLP) network with three fully connected layers for classification. To ensure sufficient training and prevent overfitting, we apply dropout with a rate of 0.5 in the MLP and use early stopping with a patience of 50 epochs. This means that if the validation loss does not decrease for 50 epochs, training will be halted. Other hyperparameters include a learning rate of 0.005, a weight decay of 0.0001, and a batch size of 8 graphs. The Adam algorithm, with its default parameters, is used as the optimizer, and cross-entropy loss is employed to compute the difference between the real and predicted labels.

### D. Metastasis Probability Prediction Explainer

GNNExplainer [24] generates interpretations for arbitrary neural networks and various mining tasks from the perspective of network structure and node attributes. It aims to identify the subgraph structures most relevant to prediction results, facilitating the interpretation of these results. Specifically, the Graph Mask and Feature Mask are used to screen structures and features, respectively. These two masks are treated as trainable matrices, randomly initialized, and combined with the original graph through element-wise multiplication. This task is then framed as an optimization problem to maximize the mutual information between the original graph’s prediction and the interpretation graph. Assuming that the selected structures and features that are most relevant to the model prediction results Y are 𝒢_*s*_ and **X**_*s*_ respectively, then their importance can be measured by the mutual information MI:

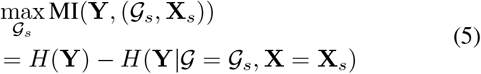

The entropy *H*(𝕐) is constant because the GCN is already trained, so the maximum mutual information between the prediction 𝕐 and the explanation (𝒢_*s*_, **X**_*s*_) is equal to the minimum conditional entropy *H*(𝕐|*𝒢* = 𝒢_*s*_, **X** = **X**_*s*_), which can be expressed as:

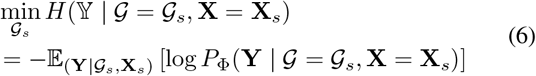

If we treat𝒢_*s*_ ∼ *𝒢* as a random graph variable, the objective in Eq.6 becomes:

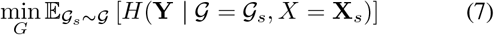

This can be given by Jensen’s inequality with convexity assumption, as the following upper bound:

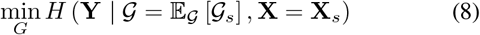

The conditional entropy in Eq.6 can be optimized by replacing the 𝕐_*G*_ [𝒢 _*s*_] with a masking of the computation graph of the adjacency matrix *A*_*c*_ ⊙ *σ*(*M*). Here, *M* ∈ ℝ^*n×n*^ and *F* ∈ℝ^*n×*1^ are learnable masks, ⊙ is the element-wise product, and *σ* denotes the sigmoid function that maps the mask to the range [0, 1].

In some applications, users care more about how to use trained model predict wanted labels, we can modify the conditional entropy objective in Eq.8 with a cross-entropy objective between the label class and the model prediction:

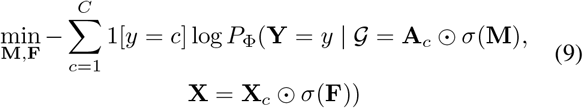

GNNExplainer is initialized with a trained graph model and individual graph data. For each graph, GNNExplainer is trained for 200 epochs to generate its unique feature mask and graph mask. To obtain a general explanation for the entire dataset, we determine the importance score of individual nodes by summing the scores of all input edges of a node and averaging by node degree. The node importance scores are calculated separately for the metastatic and non-metastatic classes.

### E. Data Partitioning and Robustness Assessment

When training the classification models, we used a stratified K-fold cross-validation procedure to divide the dataset. The dataset was randomly split into five folds, each containing an equal number of positive and negative samples. One fold was used as the testing dataset, and the remaining four folds were used as the training dataset, with half of the testing dataset reserved for validation. This resulted in 80% of the data being used for training and 10% each for validation and testing. For each split, we conducted five independent repeated experiments, and the evaluation indicators were averaged from the performances of a total of 25 models. Before training the explainer model, we retrained the GCN model with the newly split dataset. The dataset was randomly divided into three parts: 80% for training, 10% for validation, and 10% for testing. We used the same training strategy and hyperparameters as those used in the classification model experiments. The newly trained GCN model and the entire dataset were then input into the GNNExplainer model to generate the feature mask and graph mask.

## III. RESULTS AND DISCUSSION

### A. Precisely Metastasis Probability Prediction GCN Tool

In classification tasks, prediction results can be categorized into four classes: true positive (TP), false positive (FP), false negative (FN), and true negative (TN). Accuracy denotes the percentage of testing samples correctly classified by the classifier and is defined as:

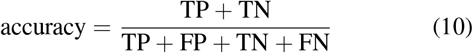

The F1 score measures the accuracy of binary classification models, considering both precision and recall. The F1 score is the harmonic mean of precision and recall, with a maximum value of 1 and a minimum value of 0. It is defined as:

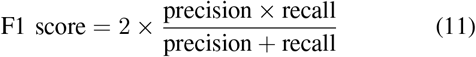

Where precision 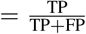, recall 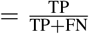.

To explore whether graph structures aid in the detection of cancer metastasis, we compared Graph Convolutional Networks (GCNs) and graph transformers with structure-agnostic models, such as Multi-Layer Perceptrons (MLPs) and transformers (Fig. 2). All four models use the same architecture shown in Fig.1. When GCNs ignore the normalized Laplacian, they degrade into MLP models, resulting in GCNs and MLPs having the same number of parameters. Compared to transformers, graph transformers calculate attention scores between nodes and their neighbors instead of all other nodes, leading to the same number of trainable parameters for both models. Table 1 shows the statistics of each model’s parameters. The experimental results are shown in Table 2. GCNs perform the best, achieving approximately 90% accuracy. MLPs achieve only 73.12% accuracy, as they consider only node features and are thus limited. Similarly, transformers fall short, with an accuracy of about 72.11%, since vanilla transformers might aggregate redundant information from distant nodes. Graph transformers address this by calculating attention scores with neighboring nodes only, achieving better accuracy at about 86%. Overall, incorporating proper graph structures can significantly improve classification accuracy.

**TABLE II:** PARAMETERS IN EMBEDDING LAYERS.

| Models | Main layers | Parameters | Summary of layer-wise parameters | Model total parameters |
| --- | --- | --- | --- | --- |
| GCN_Net/<br>MLP | GCN layer 1/<br>Full connected layer 1 | Weight matrix: $W^{(1)} \in \mathbb{R}^{d_1 \times d_2}$,<br>$d_1 = 3, d_2 = 64$<br>Bias vector: $b^{(1)} \in \mathbb{R}^{b_1 \times 1}$, $b_1 = 64$ | $d_1 \times d_2 = 192$<br>$b_1 = 64$ | 4416 |
| | GCN layer 2/<br>Full connected layer 2 | Weight matrix: $W^{(2)} \in \mathbb{R}^{d_3 \times d_4}$,<br>$d_3 = 64, d_4 = 64$<br>Bias vector: $b^{(1)} \in \mathbb{R}^{b_2 \times 1}$, $b_2 = 64$ | $d_3 \times d_4 = 4096$<br>$b_2 = 64$ | |
| Transformer/<br>Graph Transformer | Linear transformer layer 1<br>($Q^{(1)}, K^{(1)}, V^{(1)}$) | Weight matrix: $(Q^{(1)}, K^{(1)}, V^{(1)}) \in \mathbb{R}^{d_1 \times d_2}$,<br>$d_1 = 3, d_2 = 64$<br>Bias vector: $(Q_b^{(1)}, K_b^{(1)}, V_b^{(1)}) \in \mathbb{R}^{b_1 \times 1}$, $b_1 = 64$ | $d_1 \times d_2 \times 3 = 576$<br>$b_1 \times 3 = 256$ | 13376 |
| | Feature aggregate layer 1 ($O^{(1)}$) | Weight matrix: $O^{(1)} \in \mathbb{R}^{d_3 \times d_4}$, $d_3 = 64, d_4 = 64$ | $d_3 \times d_4 = 4096$ | |
| | Linear transformer layer 2<br>($Q^{(2)}, K^{(2)}, V^{(2)}$) | Weight matrix: $(Q^{(2)}, K^{(2)}, V^{(2)}) \in \mathbb{R}^{d_5 \times d_6}$,<br>$d_5 = 64, d_6 = 64$<br>Bias vector: $(Q_b^{(2)}, K_b^{(2)}, V_b^{(2)}) \in \mathbb{R}^{b_2 \times 1}$, $b_2 = 64$ | $d_5 \times d_6 = 4096$<br>$b_2 \times 3 = 256$ | |
| | Feature aggregate layer 2 ($O^{(2)}$) | Weight matrix: $O^{(2)} \in \mathbb{R}^{d_7 \times d_8}$, $d_7 = 64, d_8 = 64$ | $d_7 \times d_8 = 4096$ | |

**Fig. 2:**
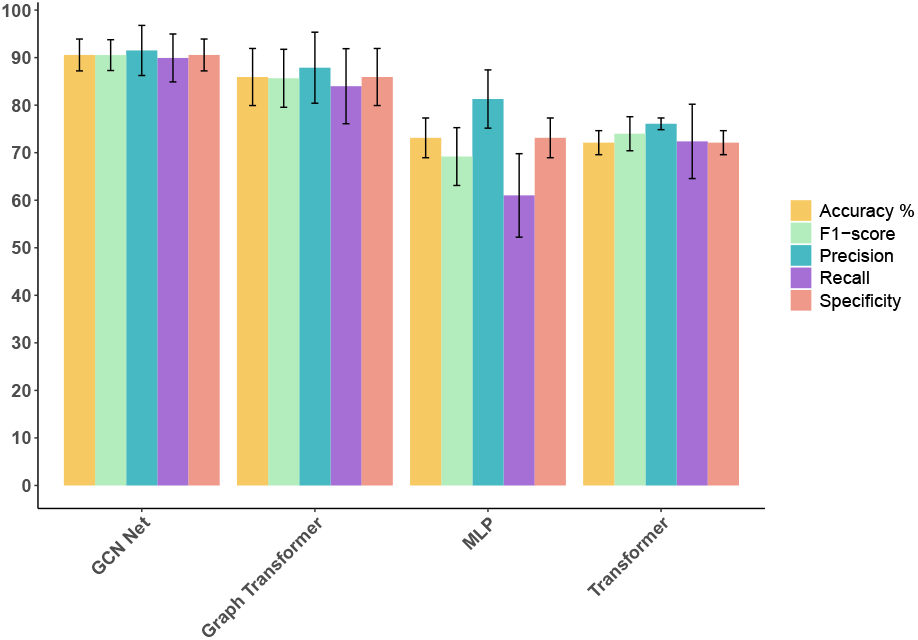
Performance of four embedding models.

### B. Multi-omics Integration Prediction Enhanced Precision in Diversity

To demonstrate that multi-omics data can improve performance, we conducted ablation experiments using GCN. The results comparing single-omics data and multi-omics data are shown in Fig. 3A, which presents the performance scores when testing the GCN model. In the single-omics data experiments, methylation (meth) achieves the highest accuracy at approximately 88%, while mutation performs the worst, with an accuracy of 77%. These results indicate that GCN achieves superior performance with multi-omics data compared to using any single omics data.

**Fig. 3:**
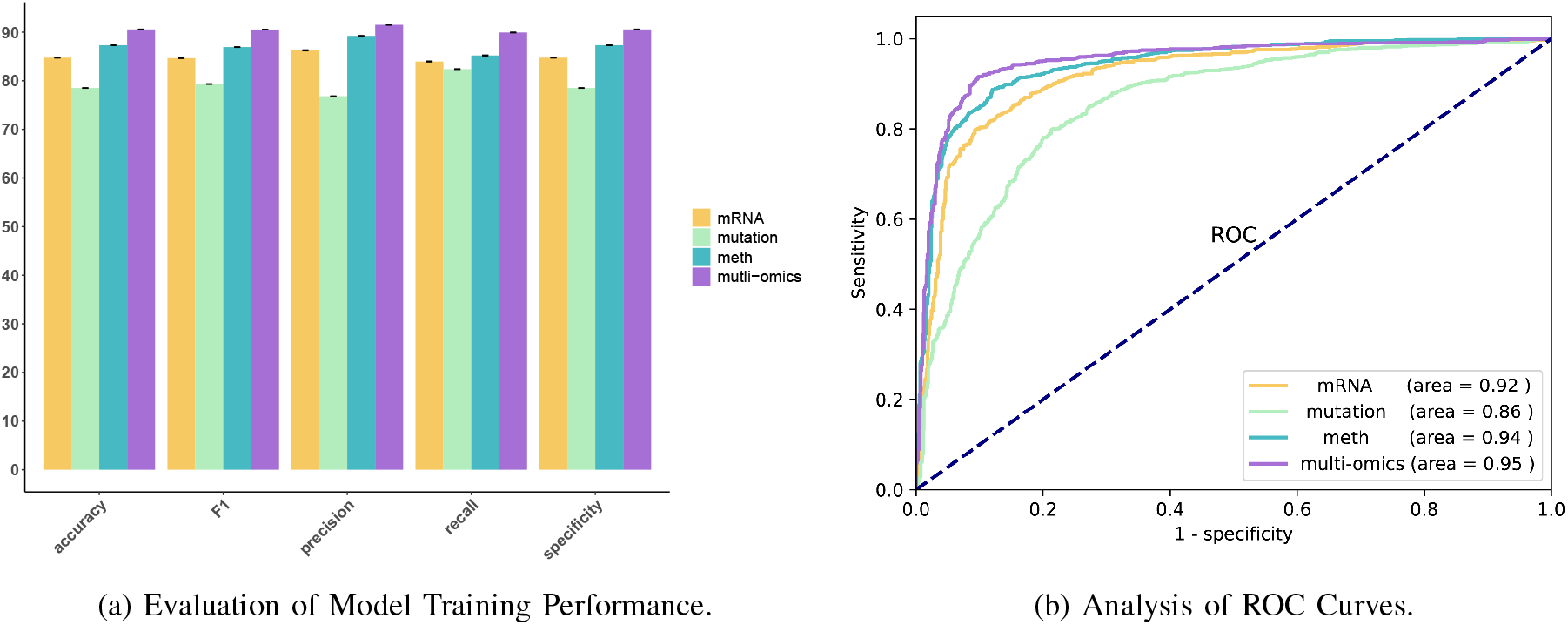
Accuracy and ROC comparison between multi-omics and single omic data. The model trained on multi-omics data outperforms those trained on single-omics data. This indicates that integrating multi-omics information enhances generalization ability and higher classification accuracy compared to models based on individual omics data.

The Receiver Operating Characteristic (ROC) curve reflects the relationship between sensitivity and specificity. The abscissa represents 1-specificity, also known as the false positive rate (FPR). The closer the X-axis is to zero, the higher the accuracy. The ordinate represents sensitivity, also known as the true positive rate, with a higher Y-axis indicating better accuracy. The Area Under the Curve (AUC) is used to measure the accuracy. A higher AUC value indicates a larger area under the curve, signifying better prediction accuracy. The closer the curve is to the top-left corner (with a smaller X and a larger Y), the better the prediction accuracy. Fig. 3B illustrates the ROC curve and AUC for both single-omics and multi-omics data. Among the single-omics data, mutation achieves the lowest AUC at 0.86. mRNA ranks second with an AUC of 0.92, and methylation performs best with an AUC of 0.94. Impressively, multi-omics data improves performance, achieving an AUC of 0.95.

### C. Interpretation method combined with MetaGXplore provides insight into metastasis pivotal genes

Utilizing the graph mask and feature mask methodologies from GNNExplainer, genes underwent ranking based on their weights, indicative of their significance within the predictive framework. Subsequently, we identified the top 500 genes for both metastatic and non-metastatic patient prognostication, facilitating further scrutiny. Fig. 4. illustrates the molecular function and cell component pathway annotation results for the top 500 genes associated with metastasis and non-metastasis patients.

**Fig. 4:**
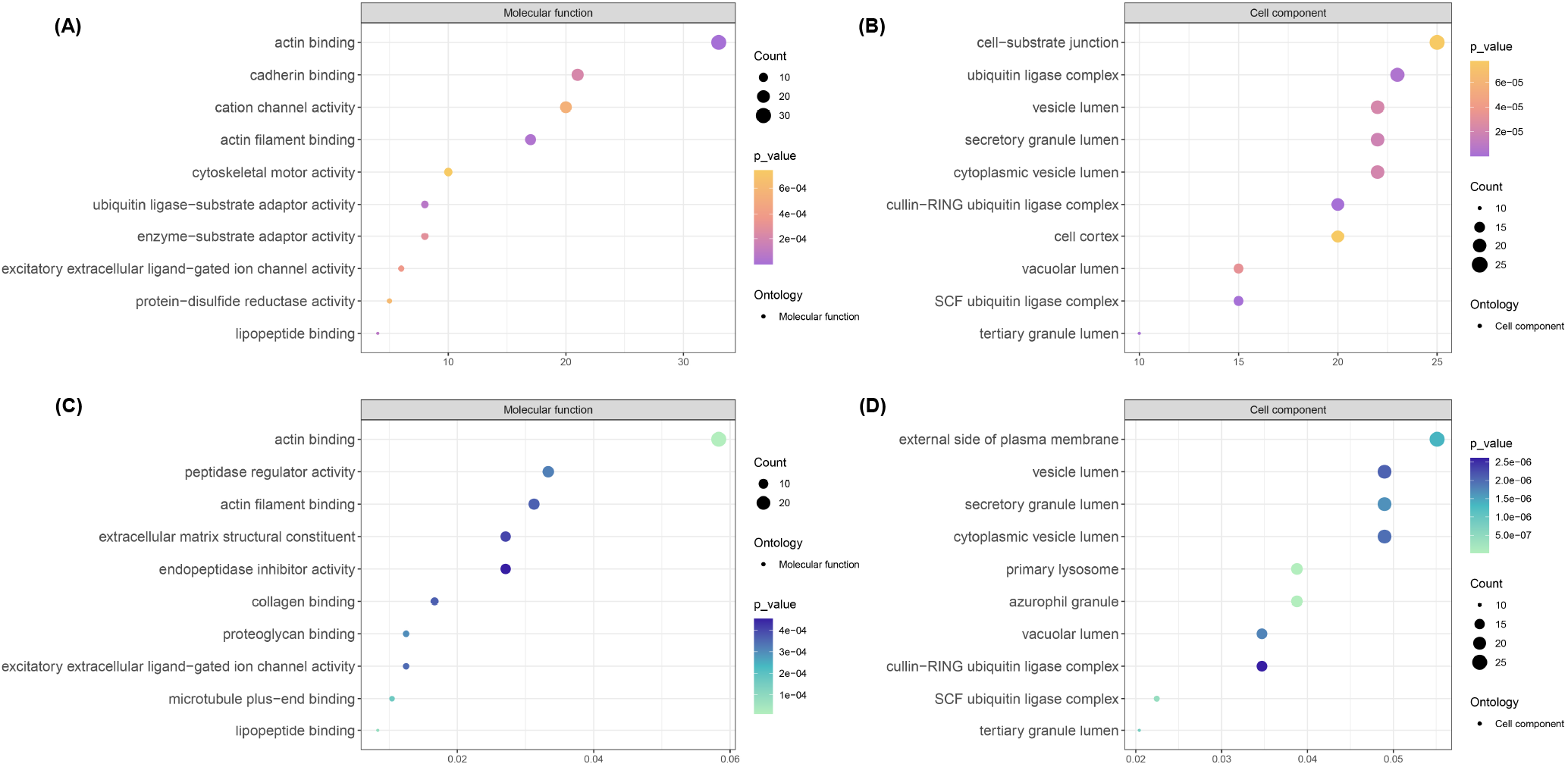
The molecular function and cell component pathway annotation result of the top 500 genes associated with metastasis (A, B) and nonmetastasis (C, D) patients.

Functional pathway annotation of gene sets highly associated with metastasis revealed their involvement across diverse biological processes, including RNA splicing, actin filament organization, anatomical structure homeostasis, cell killing, and regulation of RNA splicing (Fig. 5). These processes frequently intersect with cancer metastasis mechanisms, including cell migration, invasion, and adaptation to new environments. Additionally, we presented a network visualization delineating relevant Gene Ontology (GO) terms, with larger nodes denoting a greater number of associated genes (Fig. 5).

**Fig. 5:**
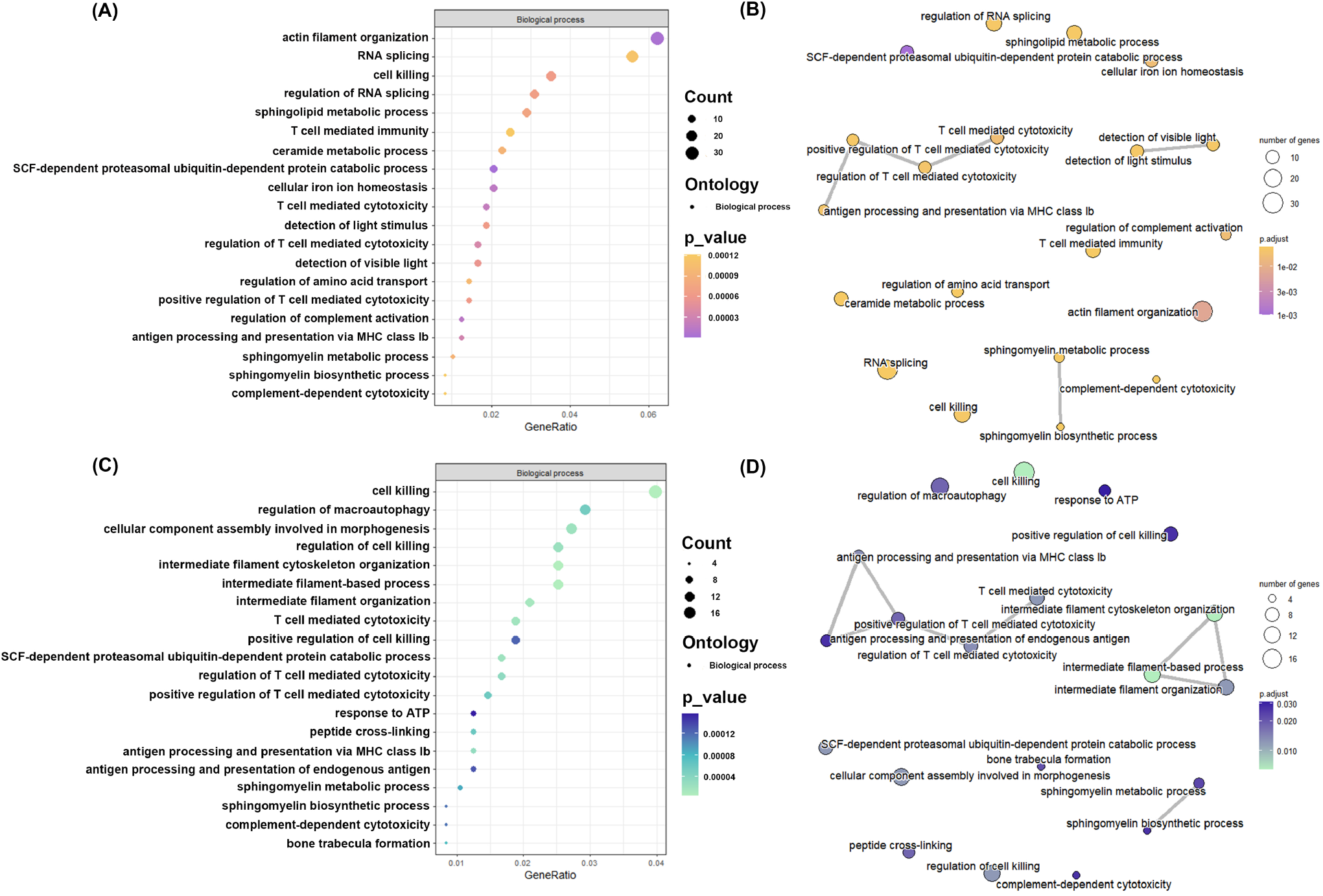
Enrichment analysis of the pivotal genes. (A) The top 500 genes associated with metastasis interpreted from MetaGXplore were enriched for biological functional pathways, showing the top twenty significantly enriched pathways. (B) Relationship between pathways of metastasis prediction-related genes. (C-D) The top 500 genes related to non-metastasis prediction were enriched in biological pathways and the relationships between pathways.

**Fig. 6:**
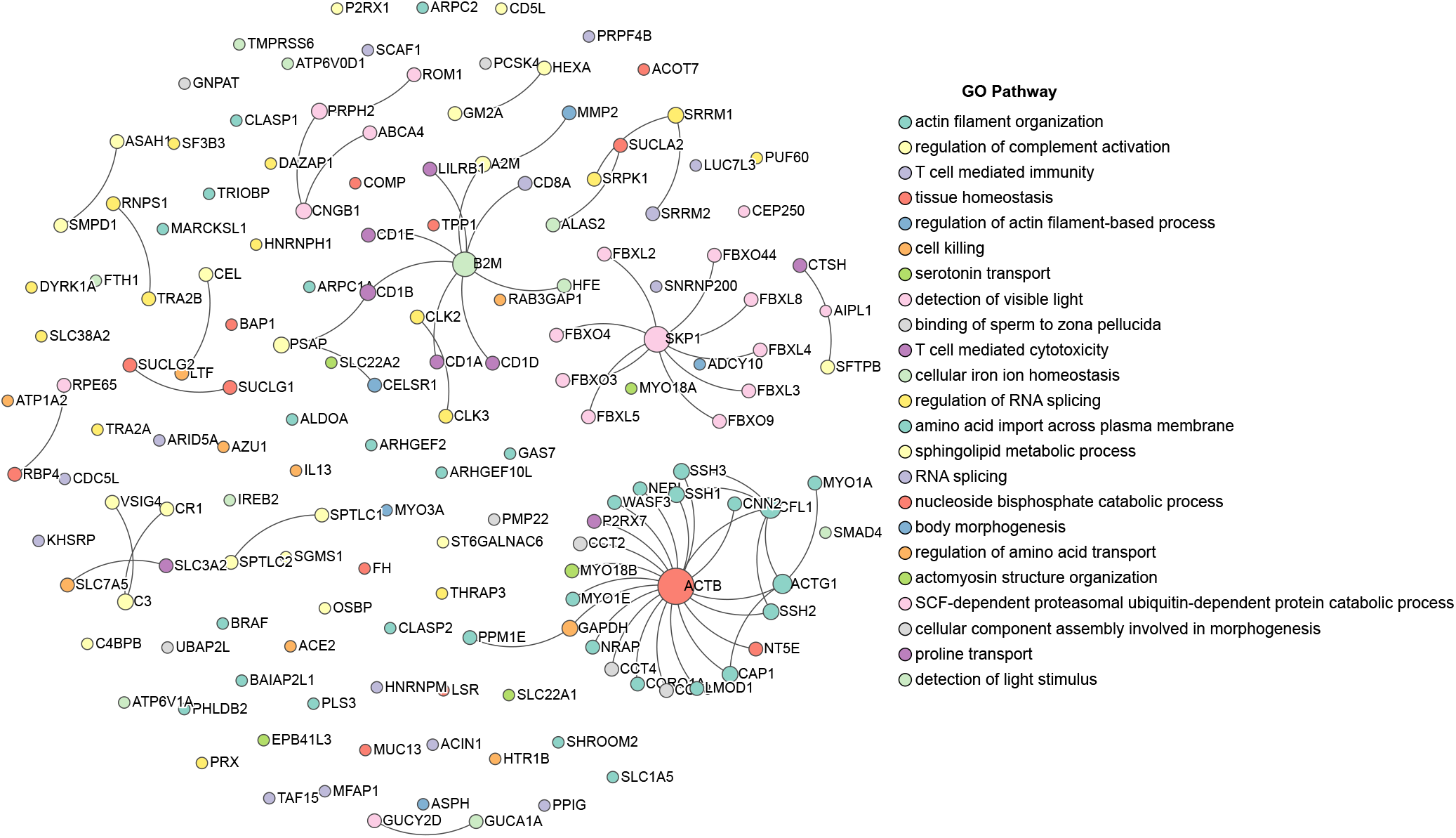
The pivotal genes associated with metastasis. The annotated sections are overlaid onto a Protein-Protein Interaction (PPI) network to examine their interrelationships, forming the largest network cluster. The colors and sizes represent the biological function annotated in GO pathways and the contribution of nodes to the network

Noteworthy pathways highlighted include T cell-mediated cytotoxicity, antigen processing, presentation via MHC class 1b, as well as various metabolic processes like sphingomyelin, sphingoglycolipid, and ceramide metabolism. This underscores the enrichment of the analyzed gene sets in biological processes and pathways pertinent to metastasis, such as cell migration, immune evasion, and metabolic reprogramming. Notably, several genes exhibited significant enrichment in RNA splicing, with a notable representation in this pathway. Among these, spliceosome components SF3B3, PRPF4B, SRRM1, and SRRM2 form a dynamic ribonucleoprotein complex assembled from small nuclear RNA (snRNA) and diverse proteins within the eukaryotic cell nucleus. Somatic mutations in pre-mRNA splicing factors represent frequent acquired mutations and early genetic events in myelodysplastic syndromes (MDS) and related myeloid tumors, alongside some solid tumors [25, 26, 27].

Through the pivotal genes linked to metastasis, we constructed a protein-protein interaction (PPI) network involving the top 500 genes, excluding singleton nodes. These genes play crucial roles in development, cell fate determination, and have been implicated in various pathological conditions. Notably, four hub genes—ACTB, B2M, SKP1, and CFL1—emerged prominently with diverse function pathways (Fig.6). The network analysis reveals a complex interplay of genes, with several hub nodes emerging as potentially critical regulators. ACTB (Actin Beta) encodes a major component of the cytoskeleton, essential for cell structure, motility, and integrity, and is often used as a reference gene in gene expression studies. B2M (Beta-2-Microglobulin) is a component of the major histocompatibility complex (MHC) class I molecules, playing a crucial role in immune system function and often studied in the context of immune-related disorders and cancer. SKP1 (S-Phase Kinase-Associated Protein 1) is a key component of the SCF (SKP1-Cullin-F-box) complex, which is involved in ubiquitin-mediated proteolysis, critical for cell cycle regulation and tumor suppression. CFL1 (Cofilin 1) is an actin-binding protein that regulates actin dynamics, crucial for cell movement, division, and is implicated in cancer metastasis and neurological disorders. Given their roles in cell structure (ACTB), immune function (B2M), cell cycle regulation (SKP1), and actin dynamics (CFL1), these genes are either already implicated in tumor metastasis or represent promising candidates for further research as potential biomarkers and therapeutic targets in cancer metastasis.

Conversely, genes bearing high weights associated with non-metastatic patients, as discerned through the interpretability analysis of MetaGXplore, revealed significant enrichment in novel pathways (Fig. **??** and **??**). Compared to counterparts at high metastatic risk, gene sets enriched in non-metastatic patients manifested fresh biological pathways, including regulation of macroautophagy, cellular component assembly, intermediate filament cytoskeleton organization, and positive regulation of cell killing. These pathways may serve to impede or deter metastatic progression by mechanisms such as bolstering tumor cell self-clearance, sustaining cellular structural integrity, and augmenting anti-tumor immune responses. This discovery furnishes a theoretical foundation for elucidating the molecular orchestration of metastasis and advancing novel anti-metastatic targeted therapies.

## IV. CONCLUSION

In this research endeavor, we have devised a pioneering deep learning model aimed at forecasting the likelihood of metastasis occurrence in patients afflicted with various cancers. Employing an interpretable framework, we have discerned a pivotal set of genes whose role in prediction is pronounced and possibly intertwined with tumor metastasis. Our findings offer a valuable resource for healthcare practitioners, furnishing them with a predictive tool for gauging the metastatic risk on a per-patient basis. Furthermore, the elucidation of this gene set lends insights into the underlying biological mechanisms governing metastasis, enriching our understanding of this intricate process.

## DATA AVAILABLE STATEMENT

Publicly available datasets were analyzed in this study. This data can be found here: The 754 cancer samples were obtained from the The Cancer Genome Atlas (TCGA; https://www.cancer.gov/). Access data and model for this article are listed in https://github.com/TJiangBio/MetaGXplore.

